# Geographic mosaic of selection by avian predators on hindwing warning colour in a polymorphic aposematic moth

**DOI:** 10.1101/2020.04.08.032078

**Authors:** Katja Rönkä, Janne K. Valkonen, Ossi Nokelainen, Bibiana Rojas, Swanne Gordon, Emily Burdfield-Steel, Johanna Mappes

**Affiliations:** University of Jyväskylä, Department of Biological and Environmental Science, Jyväskylä, Finland; Helsinki Institute of Life Sciences, University of Helsinki, Helsinki, Finland; Organismal and Evolutionary Biology Research Programme, Faculty of Biological and Environmental Sciences, University of Helsinki; Department of Biology, Washington University in St. Louis, St. Louis, Missouri, United States of America; Institute for Biodiversity and Ecosystem Dynamics, University of Amsterdam, Amsterdam, The Netherlands

**Keywords:** frequency-dependent selection, aposematism, colour polymorphism, signal variation, signal convergence, predators, predator-prey interactions, wood tiger moth, *Arctia plantaginis*

## Abstract

Warning signals are predicted to develop signal monomorphism via positive frequency-dependent selection (+FDS) albeit many aposematic systems exhibit signal polymorphism. To understand this mismatch, we conducted a large-scale predation experiment in four locations, among which the frequencies of hindwing warning coloration of aposematic *Arctia plantaginis* differ. Here we show that selection by avian predators on warning colour is predicted by local morph frequency and predator community composition. We found +FDS to be strongest in monomorphic Scotland, and in contrast, lowest in polymorphic Finland, where different predators favour different male morphs. +FDS was also found in Georgia, where the predator community was the least diverse, whereas in the most diverse avian community in Estonia, hardly any models were attacked. Our results support the idea that spatial variation in predator and prey communities alters the strength or direction of selection on warning signals, thus facilitating a geographic mosaic of selection.

## Introduction

The survival strategy of aposematism, wherein prey use warning signals that predators learn to associate with their unprofitability and subsequently avoid, has stimulated biological studies for centuries (Ruxton *et al.* 2018; Skelhorn *et al.* 2016; Merrill *et al.* 2015; Mappes *et al.* 2005; Cott 1940; Poulton 1890; Wallace 1867). In aposematism, prey benefit from lowered costs of predator education by carrying a common signal, while predators reduce risks by not attacking defended prey. This results in selection for local similarity in warning signals, a view that has been corroborated by theoretical approaches (e.g. Aubier & Sherratt 2015; Sherratt 2008; Mallet & Joron 1999; Müller 1878), laboratory experiments (e.g. Rowland *et al.* 2007; Lindström *et al.* 2001; Greenwood *et al.* 1989), and field studies (e.g. Dell’aglio *et al.* 2016; Chouteau *et al.* 2016; Borer *et al.* 2010; Kapan 2001; Mallet & Barton 1989). Nevertheless, phenotypic variation and polymorphism in aposematic organisms are widespread in nature (e.g. frogs: Rojas 2017; Siddiqi *et al.* 2004; newts: Beukema *et al.* 2016; Mochida 2011; butterflies: Merrill *et al.* 2015; moths: Brakefield & Liebert 1985; bumblebees: Plowright & Owen 1980; beetles: Bocek & Bocak 2016; Brakefield 1985; locusts: Nabours 1929; myriapods: Marek & Bond 2009; nudibranchs: Winters *et al.* 2017), which requires an evolutionary explanation.

Given that the association between prey warning signal and defence should be learned by each generation of predators (Mappes *et al.* 2014), the benefit of signal sharing depends on how often predators encounter the signal. The encounter rate then depends on both the frequency (Heino *et al.* 1998; Müller 1879) and density (Endler & Rojas 2009; Rowland *et al.* 2007; Sword 1999; Müller 1879) of prey carrying the signal. Thus, it is expected that selection on aposematism is positively frequency-dependent (+FDS), with predators avoiding the most common warning signal in a locality (Ruxton *et al.* 2018; Chouteau *et al.* 2016; Comeault & Noonan 2011; Chouteau & Angers 2011; Sherratt 2008).

On the other hand, several mechanisms have been proposed to counterbalance selection for signal monomorphism and facilitate warning colour polymorphism (reviewed in Briolat *et al.* 2018). For example, temporally and spatially varying interspecific interactions can result in geographically variable patterns of polymorphism (McLean & Stuart-Fox 2014), particularly when coupled with limited amounts of gene flow between differentially selected populations (e.g. Gordon *et al.* 2015; Aubier & Sherratt 2015; Merilaita 2001). Often these mechanisms are thought to act simultaneously, or alternate in time or space (Stevens & Ruxton 2012; Gray & McKinnon 2007; Mallet & Joron 1999) creating a geographic mosaic of selection (Thompson 2005). Although both theoretical (e.g. Holmes *et al.* 2017; Gordon *et al.* 2015; Aubier & Sherratt 2015) and experimental work (e.g. Aluthwattha *et al.* 2017; Willink *et al.* 2014) have identified several mechanisms that allow multiple morphs to persist, there is no conclusive evidence from the field and the relative importance of different selective agents is not well understood (Chouteau *et al.* 2016; Stevens & Ruxton 2012). Alas, there is little empirical evidence as to the role of predator communities on local or global morph frequencies of aposematic prey.

The variation in the degree of warning colour polymorphism shown by the wood tiger moth (*Arctia plantaginis*) across the western Palearctic provides an excellent system to study how warning signal variation is maintained in the wild (Hegna *et al.* 2015). At a local scale, predator community structure (Nokelainen *et al.* 2014) and sexual selection (Gordon *et al.* 2015; Nokelainen *et al.* 2012) have been found to alter the direction of selection on white and yellow male morphs, but no previous studies have addressed selection on a wide geographic scale and including *A. plantaginis* females, which are commonly red. We exposed artificial moths representing the three hindwing colour morphs (white, yellow, red), to local predators in a field experiment spanning across four countries, while monitoring the abundance and community structure of local predator species. We tested whether 1) selection by predators favours the locally common morph; 2) the community structure of avian predators is associated with the predation pressure on different morphs; and 3) there is variation in the direction or strength of selection among populations, matching the local morph frequencies. Variable selection pressure is one of the main candidate mechanisms for the maintenance of polymorphism. By our work, we provide the best-documented case to date of a geographic mosaic of selection on warning signals at broad spatial scales.

## Material and methods

### Study system

Adult wood tiger moths, *Arctia plantaginis* (Erebidae: Arctiinae; formerly *Parasemia;* see Rönkä *et al.* 2016 for classification), show conspicuous warning colours and possess a chemical defence fluid, which contains pyrazines (Burdfield-Steel *et al.* 2018b; Rojas *et al.* 2017) and is a deterrent to avian predators (Burdfield-Steel *et al.* 2018a; Rojas *et al.* 2017). Their warning coloration varies throughout their Holarctic distribution, but local polymorphism is common too (Hegna *et al.* 2015). In the western Palearctic male hindwing colour is either white or yellow, or varies more continuously between yellow and red as seen in females. We selected four study locations that represent the colour variation continuum from monomorphic to polymorphic *Arctia plantaginis* populations in the western Palearctic (Figure 1). For the purposes of this study, we consider both sexes to belong to the white, yellow or red morph based on their hindwing colour, and simplify the continuous hindwing coloration of females and Georgian males into two classes: yellow and red (categorized by human eye in 6 grades as in Lindstedt *et al.* 2011), here grades 1-2 are determined yellow and 3-6 red). Accordingly, Scotland is monomorphic with yellow males and females, Georgia is mostly red with 4.3% of males being yellow, and Estonia and Finland are polymorphic with all females caught between 2013-2015 classified as red, and males as either white or yellow (Figure 1).

**Figure 1.**
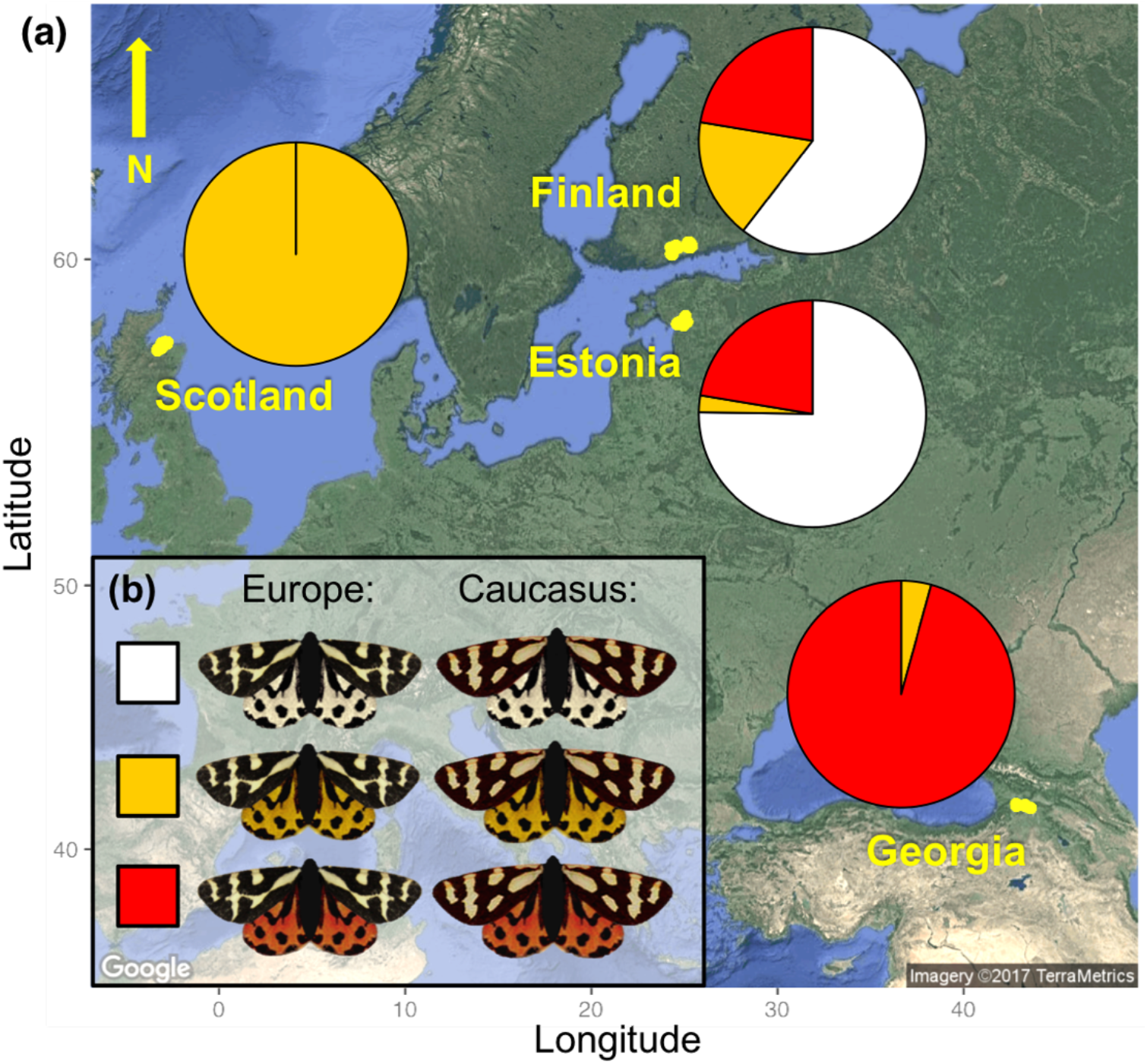
(a) Study populations and local morph frequencies and (b) moth models used in Europe and Georgia representing a local forewing pattern and white, yellow or red hindwing morph. At the monomorphic end of the monomorphic-polymorphic continuum in Scotland, both sexes have yellow hindwing coloration. In Georgia red is the dominating hindwing colour, but male coloration varies continuously towards yellow. In Estonia, white hindwing colour dominates, as the males are almost exclusively white and females red. In Southern Finland all three colour morphs are present (white and yellow male morphs and females vary continuously from yellow to red).

Wood tiger moths are widespread but often low in numbers. Therefore, colour morph frequencies were calculated by population based on annual surveys using pheromone traps and netting between 2009 and 2014 in Scotland, and 2013-2015 in Georgia, Estonia and Finland. Morph frequencies for white and yellow males, and yellow or red males in Georgia, were calculated as the average frequencies from all data available. Morph frequencies for yellow and red females were based on netting data only, as the pheromone traps only lure male moths. Because our dataset was thus biased towards male moths, we corrected the morph frequencies according to a sex ratio of 45 females to 156 males, based on a mark-release-recapture study spanning two years in Central Finland (Gordon *et al*. unpublished data). This sex ratio was used, as it is likely to depict the detectability of each morph more accurately than an even 1:1 sex ratio. The higher frequency of males to females is supported by two observations: male wood tiger moths live longer and fly more actively than females, and the adult sex ratio immediately after eclosion is slightly biased towards males even in laboratory conditions (K. Suisto, personal communication). The concluding morph frequencies (Figure 1A) are consistent with museum samples (Hegna *et al.* 2015) and laboratory stocks originating from the four study populations (Central-Finland, Estonia, Scotland and Georgia).

### Predation experiment

To estimate the attack risk of white, yellow and red hindwing colour morphs by local predators in the wild we used artificial moth models, resembling real *A. plantaginis* morphs. Models with plasticine (Caran D’Ache Modela 0259.009 Black) bodies attached to printed waterproof (Rite in the Rain ©, JL Darling Corporation, Tacoma, WA, USA) paper wings were prepared following methods described in Nokelainen *et al*. (2014). Models were constructed using pictures of one white moth hindwing and two forewings, one with a typical European pattern and another with a typical Caucasian (Georgian) pattern, which were copied and assembled in GIMP 2.8.16 SOFTWARE (GNU Image manipulation program) to create six models representing the white, yellow and red morphs in Europe and Georgia (Figure 1B). A locally common forewing type was used to reduce potential novelty effect caused by the forewing pattern (Hegna & Mappes 2014). Resemblance of the artificial models to the real moths was verified by taking measurements of reflectance from the black and coloured areas of real moth wings and printed wings with a Maya2000 Pro spectrometer (Ocean Optics) using a PX-2 Pulsed Xenon Light Source (Ocean Optics) for illumination and adjusting the model colours with Gimp (2.8.16) to match the natural wing colour as closely as possible with a calibrated (HP Colour LaserJet CP2025) printer (see spectral curves of hindwing colour in Rönkä *et al.* 2018), where identical models were used). As our study focused on the hindwing coloration, all other variables such as wing size and pattern were kept constant.

We set up 60 predation transects across the four study populations (15 in each country) in open, semi-open and closed natural habitats where the wood tiger moth and its potential avian predators were known or presumed to occur. The predation transects were set at least 500 m apart to avoid birds having overlapping territories between the transects. Along each 900 m transect 20 white, 20 yellow and 20 red artificial moth models were set individually every 15 meters using a randomized block design, so that two models of the same colour would never be next to each other. Models were pinned directly on natural vegetation, either to green leaves large enough to support their weight, or to tree trunks, as visibly as possible. All models were left in the field for a maximum of 6 days (2-6 days, 4 days on average), during the *A. plantaginis* flight season in 2014 (May 31st – July 6th in Estonia, May 26th – July 6th in Finland, June 15th – July 30th in Scotland and July 12th – August 3rd in Georgia). Predation events were recorded every 24 hours except for days of heavy rain (as birds were likely not active). For practical reasons (i.e. accessibility of mountain roads and weather conditions) the protocol was modified in Georgia. The 20 white, 20 yellow and 20 red models were set every 10 m totalling up to 600 m, left in the field for 3 consecutive days (72 h), and checked only once.

Attacks were recorded based on imprints on the plasticine body and fractures in the wings (see Supplemental Experimental Procedures). Only clear avian attacks were included in the analyses (Supplemental Table 1). Missing and attacked models were replaced with a new model of the same colour to ensure constant morph frequency during the experiment. Excluding or keeping consecutive attacks on the replaced models in the analyses did not markedly change the outcome, reported here (Table 1) for the dataset including replaced models (4004 observations) and for the dataset including original models only (3600 observations; in Supplemental Table 2). Therefore, we kept the replaced models in for all of the analyses, as it increased the sample size.

**Table 1.** Positive frequency-dependency of the estimated attack risk. (a) The model including only main effects of morph frequency and morph colour (underlined) was selected because the reduction in AIC value compared to a full model with both main effects and interaction between morph frequency and colour is higher than 2 (Δ AIC = 3.4). (b) Estimates of the best-fit model (Model 1a). Values of significance level <0.05 are bolded. Δ df denotes change in model degrees of freedom.

| (a) Model selection | $\Delta$ df | LRT | Pr(Chi) | model AIC |
| --- | --- | --- | --- | --- |
| colour * morph frequency |  |  |  | 3957.0 |
| <u>colour + morph frequency</u> | 2 | 0.1949 | 0.907 | 3953.2 |

**(b) Model 1a**
| Random effects | Variance | SD |  |  |
| --- | --- | --- | --- | --- |
| transect within country | 0.3315 | 0.5758 |  |  |
| country | 0.1633 | 0.4041 |  |  |
| Fixed effects | Estimate | SE | Z-value | p-value |
| (Intercept): colour[w] | -3.0433 | 0.2275 | -13.376 | **<0.001** |
| colour[y] | -0.0923 | 0.0940 | -0.982 | 0.3259 |
| colour[r] | -0.0841 | 0.0925 | -0.909 | 0.3633 |
| morph frequency | -0.3728 | 0.1071 | -3.481 | **0.0005** |

### Measures of predator community

To estimate the abundances of different insect-feeding birds, which are the most likely predators of wood tiger moths, we counted birds belonging to the orders Passeriformes and Piciformes (Supplemental Table 3). These counts were done once, either before or during the predation experiment, along the predation lines using a modified transect count method (see Nokelainen *et al.* 2014). Bird species observed only in one transect (out of 60), or clearly not adapted to prey on moths (e.g. crossbills), were excluded from analysis. Observations were done within 25 m from the middle of the transect in calm weather between 6 am −1 pm, when birds were most active. Shannon-Wiener diversity index (Figure 3) was calculated using R package ‘vegan’ 2.5-6 (Oksanen *et al.* 2013).

### Statistical analyses

To investigate how local predator community affects the direction and strength of selection on wood tiger moth morphs, we constructed generalised linear mixed models. Because the artificial moths were presented to predators over a different number of days in each transect, the attack risk (attacked or not) within a day exposed was used as the response variable for all analyses, modelled with a binomial distribution and a logit link function. First, we tested whether predators select for wood tiger moth warning colours in a frequency-dependent manner across populations (Figure 2). For this, we used local morph frequency calculated from field monitoring data and its interaction with colour morph as the explanatory variables in Model 1 (Table 1). Transect ID, nested within country, was set as the random factor to account for the nested spatial structure of the study design.

**Figure 2.**
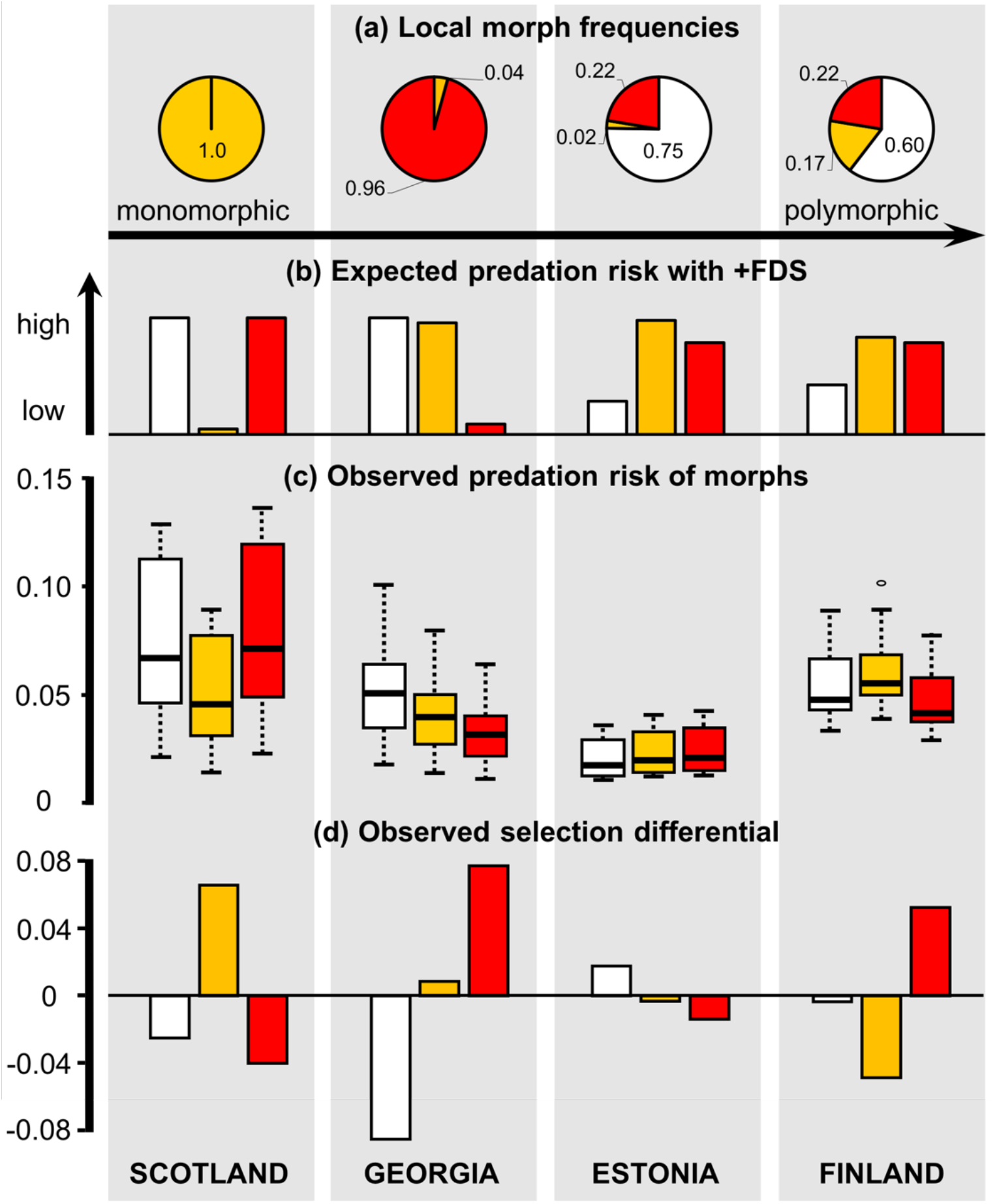
Wood tiger moth morph frequencies compared to expected and observed predation risk and selection differential by country. (a) Local morph frequencies calculated from annual monitoring data, (b) expected attack risk according to the +FDS hypothesis, where each morph is attacked according to its local frequency, (c) observed predation illustrated as GLMM estimates of daily attack risk for each morph by country and (d) observed difference in attacks per morph compared to a situation where all morphs would be attacked equally. Morph colours (white, yellow, red) as in Figure 1.

To test for predator community composition effects, the dimensions of the bird count data, consisting of 12 genera, was first reduced with a principal component analysis using the R function ‘princomp’. To avoid overparameterization, the main effects of the first three resulting components (explaining 44.7 %, 33.7 % and 8.5 % of the variation in predator community), and their three-way interactions with morph colour and country, were included one by one as explanatory variables in three separate GLMMs (Table 2). Country was included as an explanatory variable to test for local differences in selection and thus transect ID alone was set as a random effect to each model.

**Table 2.** The interaction effect of predator community and location (country, C) on the attack risk towards the wood tiger moth colour morphs. (a) Model selection starting from the main effects, interactions and a three-way interaction between principal component 1 (PC1), country (C) and colour morph (colour) as the explanatory variables, with the best-fit model underlined. (b) Estimates from the selected model (Model 2) with a significant three-way interaction of principal component 1, colour morph and country. (c) Model selection for principal component 2. Principal component 2 had no significant effects on attack risk, and thus estimates are not shown. (d) Model selection for principal component 3. Principal component 3 had no significant effects on attack risk, and thus estimates are not shown. Values of significance level <0.05 are bolded. Δ df denotes change in model degrees of freedom.

| **(a) Model selection with PC1** | **$\Delta$ df** | **LRT** | **Pr(Chi)** | **model AIC** |
| --- | --- | --- | --- | --- |
| <u>PC1*colour*C</u> |  |  |  | 3956.1 |
| PC1+colour+C+PC1:colour+PC1:C+colour:C | 6 | 14.35 | **0.026** | 3958.4 |

The asterisk (\*) denotes both main effects and interaction terms used.
| **(b) Model 2** |  |  |  |  |
| --- | --- | --- | --- | --- |
| **Random effects** | **Variance** | **SD** |  |  |
| transect | 0.2779 | 0.5272 |  |  |
| **Fixed effects** | **Estimate** | **SE** | **Z-value** | **p-value<sup>1)</sup>** |
| (Intercept): colour[w], C[Finland] | -3.556 | 0.438 | -8.126 | **<0.001** |
| colour[y] | 0.869 | 0.407 | 2.134 | **0.033** |
| colour[r] | 0.370 | 0.440 | 0.840 | 0.401 |
| PC1 | 0.139 | 0.088 | 1.590 | 0.112 |
| C[Estonia] | -0.309 | 0.621 | -0.498 | 0.619 |
| C[Georgia] | 0.618 | 0.647 | 0.955 | 0.340 |
| C[Scotland] | 0.843 | 0.477 | 1.766 | 0.077 |
| PC1 : colour[y] | -0.164 | 0.083 | -1.989 | **0.047** |
| PC1 : colour[r] | -0.115 | 0.089 | -1.299 | 0.194 |
| C[Estonia] : colour[y] | -0.463 | 0.606 | -0.765 | 0.444 |
| C[Estonia] : colour[r] | 0.297 | 0.618 | 0.481 | 0.631 |
| C[Georgia] : colour[y] | -0.773 | 0.592 | -1.306 | 0.192 |
| C[Georgia] : colour[r] | -0.041 | 0.616 | -0.067 | 0.947 |
| C[Scotland] : colour[y] | -1.169 | 0.445 | -2.624 | **0.009** |
| C[Scotland] : colour[r] | -0.345 | 0.472 | -0.730 | 0.466 |
| PC1 : C[Estonia] | -0.123 | 0.097 | -1.273 | 0.203 |
| PC1 : C[Georgia] | -0.143 | 0.111 | -1.280 | 0.201 |
| PC1 : C[Scotland] | -0.159 | 0.103 | -0.551 | 0.121 |
| C[Estonia] : PC1 : colour[y] | 0.197 | 0.093 | 2.115 | **0.035** |
| C[Estonia] : PC1 : colour[r] | 0.176 | 0.099 | 1.781 | **0.075** |
| C[Georgia] : PC1 : colour[y] | 0.105 | 0.105 | 0.991 | 0.322 |
| C[Georgia] : PC1 : colour[r] | -0.033 | 0.113 | -0.294 | 0.769 |
| C[Scotland] : PC1 : colour[y] | 0.256 | 0.100 | 2.550 | **0.011** |
| C[Scotland] : PC1 : colour[r] | 0.091 | 0.102 | 0.892 | 0.372 |

| **(c) Model selection with PC2** | **Δ df** | **LRT** | **Pr(Chi)** | **model AIC** |
| --- | --- | --- | --- | --- |
| PC2*colour*country |  |  |  | 3962.6 |
| PC2+colour+C+PC2:colour+PC2:C+colour:C | 6 | 4.413 | 0.621 | 3955.0 |
| <u>PC2+colour+C+PC2:colour+colour:C</u> | 3 | 3.334 | 0.343 | 3952.4 |
| PC2+colour+C+colour:C | 2 | 3.923 | 0.141 | 3952.3 |
| PC2+colour+C | 6 | 16.737 | **0.010** | 3957.0 |
| colour*country | 1 | 0.545 | 0.460 | 3950.8 |

| **(d) Model selection with PC3** | **Δ df** | **LRT** | **Pr(Chi)** | **model AIC** |
| --- | --- | --- | --- | --- |
| PC3*colour*country |  |  |  | 3963.6 |
| PC3+colour+C+PC3:colour+PC3:C+colour:C | 6 | 5.190 | 0.520 | 3956.8 |
| <u>PC3+colour+C+PC3:colour+colour:C</u> | 3 | 1.400 | 0.706 | 3952.2 |
| PC3+colour+C+colour:C | 2 | 4.599 | 0.100 | 3952.8 |
| PC3+colour+C | 6 | 16.726 | **0.010** | 3957.5 |
| colour*country | 1 | 0.034 | 0.854 | 3950.8 |

Statistical models were simplified using a backward stepwise deletion method based on Akaike Information Criterion. Variables were excluded one by one from the full models and the new model was accepted if the deletion reduced the AIC value more than 2 units, until only main effects or significant interactions were left in each model. All analyses were performed with R (RCoreTeam 2013) in RStudio 0.99.491 (RStudio Team 2015), using the package *lme4* (Bates *et al.* 2015).

## Results

### Positive frequency-dependent selection

Altogether, we observed a total of 718 bird attacks on the 4004 artificial moths. The relative attack risk of each colour morph was lower when the natural frequencies of the respective morph were higher in relation to the others (Table 1, Figure 2). Also, the morphs with intermediate local frequencies show corresponding levels of attack risk (Figure 2). This effect did not depend on colour morph itself (Table 1), as expected if the local predator avoidance depends more on local morph frequency than on morph colour.

### Predator community

The attacks were not evenly distributed across countries or transects (Figure 2C). Predation pressure varied between and within countries, being highest in Scotland and lowest in Estonia (Figure 2C). Georgia had the lowest amount of insect feeding birds observed (2.1 per 100 meters) compared to Finland (2.6), Scotland (4.0) and Estonia (4.4), respectively. Georgia also had the least diverse predator community measured with Shannon-Wiener diversity index, whereas Estonia was most diverse, followed by Scotland (Figure 3). Across countries, the three most commonly observed potential predators included the common chaffinch, the willow warbler (replaced by green warbler in Georgia) and the great tit (Supplemental Table 3), the latter of which was observed to attack the artificial moths. The first three principal components (PC1, PC2 and PC3) that explained 44.7 %, 33.7 % and 8.5 %, respectively, captured 87.0 % of variance in the predator community data. PC1 was dominated by Sylvidae (warblers), Fringillidae (finches) and Muscicapidae (flycatchers), which loaded in the negative end, whereas the positive end of the axis was loaded with Paridae (tits) (Figure 3). PC2 was dominated by Paridae and PC3 with Fringillidae, Muscicapidae and Troglotydidae (the Eurasian wren) (see Supplemental Table 4 for factor loadings).

**Figure 3.**
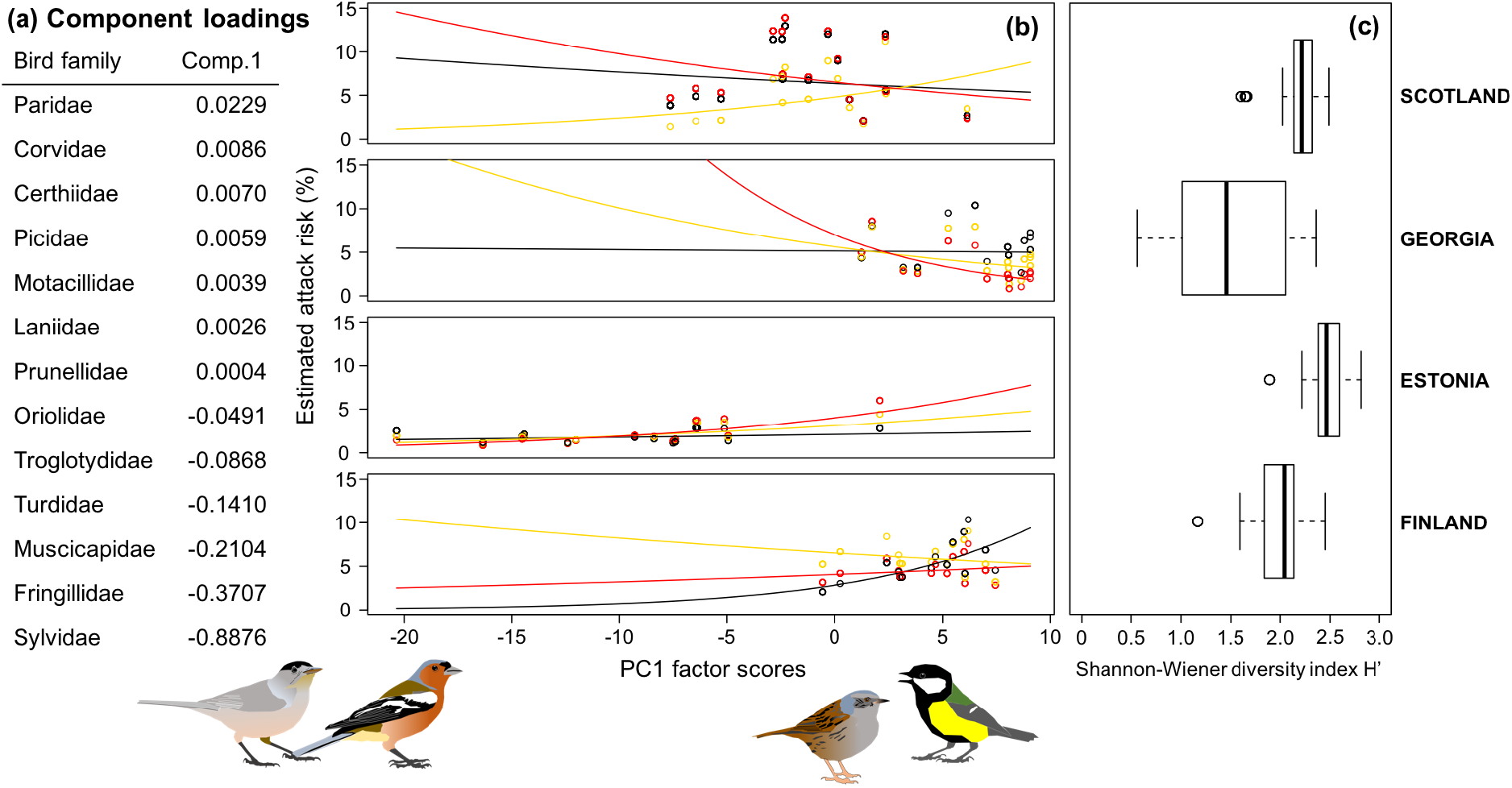
Community composition and diversity of insectivorous birds per population. Factor scores and loading of the first principal component describing 44.7% of the total variation of bird communities across countries. Panel (a) family level component loadings, panel (b) the three-way interaction effect of predator community on estimated attack risk of different colour morphs (black line corresponds to the white morph) at each transect illustrated by population, panel (c) Shannon-Wiener diversity indexes calculated per transect and plotted by population. The x-axes on both panels b and c represent factor scores for principal component 1.

### Significant association between predator community structure and selection

A consecutive analysis, where the effect of predator community on the attack risk of each moth colour morph was addressed, revealed a significant three-way-interaction between moth colour, country and PC1 (Model 2, Table 2a, Figure 3). This significant interaction means that the variation in predator community structure captured by PC1 is associated with predation pressure on different colour morphs, but the direction of the association is different between countries (i.e. between local communities). PC2 and PC3 were not significantly associated with predation pressure (Table 2C and 2D).

## Discussion

Our experiment is among the first experimental approaches integrating community-level interactions into the study of selection on warning signals (Aluthwattha *et al.* 2017; Nokelainen *et al.* 2014; Valkonen *et al.* 2012; Mochida 2011), and the first to do so on such a large geographical scale. With a wide-ranging field experiment spanning populations varying in their degree of polymorphism, we demonstrate that local bird predators avoid locally common morphs, but also that both the strength and direction of selection on warning colour varies geographically. We found that changes in local predator communities drive geographic variation in selection despite positive frequency-dependence. Local predator-prey interactions are thus contributing to the maintenance of both geographic variation and local polymorphism in warning signals.

Local avian predators appear as a key in driving warning colour evolution, which can take different evolutionary trajectories over a geographic scale. Here, the predator community had a significant, but different effect to attack risk towards each morph in different countries. The first component from the principal component analysis, explaining 44.7 % of the variation in the abundances of insectivorous birds in different families, significantly affected estimated risk of attack. However, it did so differently towards each morph in the different countries. Interpreting the component loadings and model estimates (Table 2, Figure 3), the Paridae (e.g. tits) and Prunellidae (consisting of only one species, the dunnock, *Prunella modularis*) selected for different morphs in different countries. Our results corroborate the predator community effects found by Nokelainen *et al.* (2014), as we also found that in Finland the yellow morph is favoured in communities characterised by Paridae whereas the white morph is favoured in communities characterised by Prunellidae. In contrast, an opposite effect was found in Scotland where the yellow morph dominates, suggesting that local predators can select for different colours in different populations.

Our experiment showed that across countries locally dominating colour morphs were attacked least, as predicted by +FDS. Thus, warning signal efficacy is enhanced with increasing frequency of similarly signalling individuals as predicted due to the number-dependence of predator learning and memorisation. Nonetheless, we found geographic variation in the strength of predator-induced selection. Comparison with previous experiments in those study areas that overlap (Nokelainen et al. 2014) also reveal temporal differences. We found high overall predation pressure in Scotland where the yellow morph was in favour compared to other study locations. Although Nokelainen *et al*. (2014) did not detect positive frequency dependency, they also found much higher overall attack rates in Scotland compared to Southern Finland and Estonia. On the other hand, Nokelainen *et al.* (2014) found that yellow males were significantly less attacked than white males in Southern Finland, whereas in our study the yellow morph tended to have more attacks than the other morphs. Interestingly, the frequency of yellow and white morphs varies in Southern Finland in a biannual cycle (Galarza *et al.* 2014), and the yellow morph was more common during Nokelainen *et al.’s* (2014) study, whereas in contrast the white was more common during our experiment, suggesting again that the locally most common morphs have advantage. Temporal fluctuations in local predator-prey interactions could therefore plausibly explain why estimates of predation pressure on different colour morphs conducted in different years have varied.

All morphs were attacked at equally low levels in Estonia, which implies spatial variation in the strength of selection or even locally relaxed natural selection on the warning signal. The low predation pressure is not explained by a low number of predators, as there were more insectivorous birds in Estonia than in any other study site (Supplemental Table 3). The bird community composition in Estonia differed from the other countries though, suggesting that the strength of selection was lower in diverse communities characterized by Sylvidae, Fringillidae, Muscicapidae, Turdidae, Troglotydidae and Oriolidae, as opposed to when Paridae (e.g. tits) characterized the community. Other properties of the predator community that can affect the strength of selection on warning signals include the relative abundance of naïve vs. experienced predators (Mappes *et al.* 2014), predators’ capacity to learn many different signals (Beatty *et al.* 2004), broad generalisation between the morphs (Sherratt 2008; Balogh & Leimar 2005), conflicting selection by different predators (Nokelainen *et al.* 2014; Valkonen *et al.* 2012), and the spatial arrangement of predators in relation to prey (Endler & Rojas 2009).

In temperate regions, most insectivorous birds are migratory and prey population sizes are highly variable due to interseasonal weather variability. This is likely to cause variation in the relative abundances of naïve predators across the breeding season. Furthermore, local seasonal communities are continuously changing, altering the direction and/or strength of selection on warning signals (Mappes *et al.* 2014). Siepielski *et al.* (2013) reviewed directional selection on phenotypes, and found that selection tends to vary more in strength than in direction between populations, with most of their examples coming from mid-latitudes in the northern hemisphere. Most experimental evidence of +FDS in the wild, however, comes from tropical systems (Comeault & Noonan 2011; Chouteau & Angers 2011; Mallet & Barton 1989), where the prey and predator community composition is temporally less variable (Mittelbach *et al.* 2007). In such communities, strong +FDS can lead to very accurate mimicry between warning coloured prey, whereas in more variable conditions, higher levels of variation and polymorphism can be maintained.

The paradoxical maintenance of local polymorphism despite +FDS could thus be explained by spatial and temporal variation in morph survival combined with individuals migrating between the subpopulations (Gordon *et al.* 2015; McLean & Stuart-Fox 2014; Joron *et al.* 1999). Differences in the level of population isolation, and thus gene flow between them, could explain part of the geographic variation in wood tiger moth warning colours. Population genetic evidence is both for and against this hypothesis: the red-dominated Georgian subspecies *A. p. caucasica* occuring in the mountains of Caucasus is genetically isolated to some degree from the rest of the Western Palearctic samples (Rönkä *et al.* 2016; Hegna *et al.* 2015). However, although the monomorphic yellow population in Scotland is also remote, it clusters together with the Finnish and Estonian samples based on both nuclear and mitochondrial genes (Rönkä *et al.* 2016) and microsatellites (Hegna *et al.* 2015), indicating no restrictions on gene flow. The long-term co-existence of multiple morphs and the low genetic differentiation among polymorphic populations in Finland, Estonia and the Alps with yearly variation in genetic structure (Galarza *et al.* 2014) do suggest a role for gene flow along with varying predation pressure in maintaining local populations at different frequencies.

As recently noted by several authors (e.g. Skelhorn *et al.* 2016; Chouteau *et al.* 2016; Nokelainen *et al.* 2014), more experimental work is needed to clarify predator-prey interactions at the community level in order to understand how selection is driving the evolution of warning signals in diverse natural ecosystems. Our experiment is so far the most comprehensive analysis showing how spatio-temporal variation in predator-prey communities affects the maintenance of within-species variation and evolutionary pathways to biodiversity.

## Acknowledgements

We thank Tõnis Tasane, Sally Rigg, Heli Kinnunen, Diana Abondano and Gocha Golubiani for help with the predation experiments; Otso Häärä, for counting birds in Finland; people at JYU who helped cutting out moth models; Kaisa Suisto, D. Abondano, Liisa Hämäläinen, Morgan Brain and Sini Burdillat, for rearing moths; and Jimi Kirvesoja for managing databases. The Agency of Protected Areas from the Georgian Ministry of Environment and Natural Resources Protection, provided the collection permits for Georgia; we are indebted to Khatuna Tsiklauri for help, and to the Okromelidze family for hospitality. Members of the ‘Plantaginis Journal Club’, Rose Thorogood, Øysten Opedal and two anonymous reviewers provided comments that improved an earlier version of the manuscript.

This study was funded by the Centre of Excellence in Biological Interactions (Project 284666 to JM).

The authors have no conflict of interest to declare.

## Statement of authorship

JM, KR, ON, JV, BR and SG designed the experiment. All authors contributed to the fieldwork. KR, JV and ON did the statistical analysis. KR led the writing and all authors contributed to it and accepted the final version.

## Data accessibility statement

Should the manuscript be accepted, the data supporting the results will be archived in the University of Jyväskylä data repository JYX.

