## Supplemental for "Geographic mosaic of selection by avian predators on hindwing warning colour in a polymorphic aposematic moth"

**Supplemental material**

**Supplemental Experimental Procedures**
**(related to Methods, Model 1 and Table 1 in the main text).**

*Estimated attack risk with and without potential attacks, missing models and replacement models*

Attacks were classified in the field as potential attack mark (1), clear attack mark (2), strong attack (3) and missing model (4). First, we tested whether or not the inclusion of missing moth models (values 4) and those models with markings classified as potential attacks (values 1) in the statistical analysis would change the analyses' outcomes (Table S1). Here, we used colour morph as the explanatory variable and transect ID as a random factor. Including the missing models and potential attack markings as attacks in our analyses increased model AIC values considerably in comparison to models where missing models and potential attacks were assigned as not attacked. For this reason, and to make sure that we were only counting true avian attacks, we only considered models with clear or strong beak marks as attacked in all further analyses.

Attacked and missing models were replaced with new models of the same colour to a nearby location to keep the morph frequency constant during the experiment. To test whether including or excluding attacks on the replacement models would change the analysis outcomes, we run the first model (Model 1) with and without attacks on replaced artificial moths (in the latter case, assuming that once attacked, the artificial moths died and could not be attacked again). Attacks to these replacement models are included in Model 1a (Table 1 in the main text), and not included here in Model 1b (Table S2).

**Table S1 (related to Supplemental Experimental Procedures and Methods in the main text).** Comparison of datasets including or excluding missing artificial moths and potential attack marks as attacks and with or without attacks on replaced artificial moths separately for each country. All datasets were tested with generalized linear mixed model fit by maximum likelihood (Laplace Approximation) and binomial family with a logit link function. Formula used was glmer(cbind(attack,time)~colour+(1|transect). Values of significance level <0.05 are bolded.

| Country | Markings | Repeats included | No of obs. | df | AIC | LRT | p-value |
|---|---|---|---|---|---|---|---|
| Scotland | clear | yes | 1088 | 2 | 1343.6 | 9.6804 | **0.007905** |
| | all | yes | 1195 | 2 | 1805.5 | 7.7808 | **0.02044** |
| | clear | no | 900 | 2 | 1078.6 | 5.1345 | 0.07675 |
| | all | no | 899 | 2 | 1401.5 | 4.1722 | 0.1242 |
| Georgia | clear | no | 900 | 2 | 769.0 | 6.0527 | **0.04849** |
| | all | no | 900 | 2 | 987.7 | 3.1712 | 0.2048 |
| Estonia | clear | yes | 968 | 2 | 705.9 | 0.5334 | 0.7659 |
| | all | yes | 1057 | 2 | 1278.2 | 0.9968 | 0.6075 |
| | clear | no | 900 | 2 | 664.6 | 0.3458 | 0.8412 |
| | all | no | 900 | 2 | 1126.2 | 0.2834 | 0.8679 |
| Finland | clear | yes | 1052 | 2 | 1133.2 | 2.9263 | 0.2315 |
| | all | yes | 1182 | 2 | 1659.8 | 1.4876 | 0.4753 |
| | clear | no | 900 | 2 | 1023.4 | 3.6839 | 0.1585 |
| | all | no | 900 | 2 | 1375.9 | 1.9643 | 0.3745 |

**Table S2 (related to Model 1a and Table 1 in the main text).** (a) Model selection and (b) estimates of the best-fitting model (Model 1b) with a dataset including only first attacks to each of the 60 models per transect (n=3600). Values of significance level <0.05 are bolded. Δ df denotes change in model degrees of freedom.

| **(a) Model selection** | **Δ df** | **LRT** | **Pr(Chi)** | **model AIC** |
|---|---|---|---|---|
| colour * morph frequency | | | | 3535.2 |
| <u>colour + morph frequency</u> | 2 | 0.2056 | 0.9023 | 3531.4 |

The asterisk (*) denotes both main effects and interaction terms used.

**(b) Model 1b**

| **Random effects** | **Variance** | **SD** | | |
|---|---|---|---|---|
| transect within country | 0.3022 | 0.5497 | | |
| country | 0.1337 | 0.3657 | | |
| **Fixed effects** | **Estimate** | **SE** | **Z-value** | **p-value** |
| (Intercept): colour[w] | -3.0721 | 0.2109 | -14.569 | **<0.001** |
| colour[y] | -0.0979 | 0.0982 | -0.997 | 0.3186 |
| colour[r] | -0.1316 | 0.0990 | -1.330 | 0.1836 |
| morph frequency | -0.3165 | 0.1127 | -2.808 | **0.0050** |

**Table S3 (related to Figure 3, Figure 4, Methods and Discussion in the main text).** Bird species observed within 25 meters to both sides of the transect centerline during transect counts. The table is arranged by species abundance and three most common species in each country (and in total) are bolded. Only bird species recorded to feed on insects and observed at more than one transect (out of 60) were included as potential predators in analysis (Model 4, Table 4, Figure 3, Figure 4). O=Order (Pa=Passeriformes, Co=Columbiformes, Pi=Piciformes, Ch=Charadriiformes).

| O | Family | Genus | Species | EST | FIN | GEO | SCO | Total | Incl.: | Notes |
|---|---|---|---|---|---|---|---|---|---|---|
| Pa | *Fringillidae* | *Fringilla* | *coelebs* | **102** | 19 | 7 | **66** | **194** | yes | feed insects to the young |
| Pa | *Sylvidae* | *Phylloscopus* | *trochilus* | **66** | **35** | 0 | 49 | **150** | yes | |
| Pa | *Paridae* | *Parus* | *major* | 37 | **67** | **14** | 28 | **146** | yes | |
| Pa | *Muscicapidae* | *Erithacus* | *rubecula* | 32 | 25 | 4 | **55** | 116 | yes | |
| Pa | *Sylvidae* | *Regulus* | *regulus* | 25 | 17 | 9 | 45 | 96 | yes | |
| Pa | *Troglodytidae* | *Troglodytes* | *troglodytes* | 21 | 0 | 11 | **55** | 87 | yes | |
| Pa | *Muscicapidae* | *Muscicapa* | *striata* | 29 | 15 | 8 | 24 | 76 | yes | |
| Pa | *Paridae* | *Parus* | *caeruleus* | 3 | **54** | 11 | 4 | 72 | yes | |
| Pa | *Turdidae* | *Turdus* | *merula* | 22 | 14 | 7 | 13 | 56 | yes | |
| Pa | *Paridae* | *Parus* | *ater* | 0 | 1 | 7 | 47 | 55 | yes | |
| Pa | *Sylvidae* | *Phylloscopus* | *sibilatrix* | **45** | 4 | 0 | 0 | 49 | yes | |
| Pa | *Sylvidae* | *Phylloscopus* | *collybita* | 41 | 1 | 0 | 7 | 49 | yes | |
| Pa | *Emberizidae* | *Emberiza* | *citrinella* | 0 | 18 | 0 | 23 | 41 | yes | feed insects to the young |
| Pa | *Motacillidae* | *Anthus* | *trivialis* | 21 | 12 | 0 | 5 | 38 | yes | |
| Pa | *Sylvidae* | *Sylvia* | *communis* | 25 | 0 | 0 | 11 | 36 | yes | |
| Pa | *Turdidae* | *Turdus* | *philomelos* | 15 | 7 | 0 | 10 | 32 | yes | |
| Pa | *Sylvidae* | *Sylvia* | *atricapilla* | 19 | 2 | 1 | 7 | 29 | yes | |
| Pa | *Paridae* | *Parus* | *montanus* | 11 | 13 | 0 | 4 | 28 | yes | |
| Pa | *Motacillidae* | *Anthus* | *spinoletta* | 0 | 0 | **28** | 0 | 28 | yes | |
| Pa | *Passeridae* | *Passer* | *domesticus* | 0 | 0 | 0 | 25 | 25 | no | observed in one transect only |
| Co | *Columbidae* | *Columba* | *palumbus* | 6 | 13 | 0 | 4 | 23 | no | |
| Pa | *Prunellidae* | *Prunella* | *modularis* | 3 | 7 | 2 | 11 | 23 | yes | |
| Pa | *Sylvidae* | *Sylvia* | *borin* | 20 | 2 | 0 | 0 | 22 | yes | |
| Pa | *Sylvidae* | *Phylloscopus* | *nitidus* | 0 | 0 | **22** | 0 | 22 | yes | |
| Pa | *Certhiidae* | *Certhia* | *familiaris* | 2 | 6 | 4 | 7 | 19 | yes | |
| Pa | *Fringillidae* | *Carpodacus* | *erythrinus* | 4 | 0 | 11 | 0 | 15 | yes | feed insects to the young |
| Pa | *Motacillidae* | *Motacilla* | *alba* | 6 | 1 | 3 | 2 | 12 | yes | |
| Pa | *Oriolodae* | *Oriolus* | *oriolus* | 11 | 0 | 0 | 0 | 11 | yes | feed insects to the young[1)] |
| Pa | *Fringillidae* | *Pyrrhula* | *pyrrhula* | 2 | 2 | 3 | 4 | 11 | yes | feed insects to the young |
| Pa | *Fringillidae* | *Carduelis* | *cannabina* | 0 | 0 | 5 | 6 | 11 | yes | feed insects to the young |
| Pa | *Turdidae* | *Turdus* | *viscivorus* | 7 | 0 | 0 | 1 | 8 | yes | |
| Pa | *Fringillidae* | *Loxia* | *curvirostra* | 1 | 1 | 2 | 4 | 8 | no | specialized seed eater |
| Pa | *Corvidae* | *Garrulus* | *glandarius* | 0 | 3 | 4 | 1 | 8 | yes | |
| Pa | *Fringillidae* | *Carduelis* | *carduelis* | 0 | 0 | 4 | 4 | 8 | yes | feed insects to the young |
| Pa | *Sylvidae* | *Sylvia* | *curruca* | 4 | 3 | 0 | 0 | 7 | yes | |
| Pa | *Laniidae* | *Lanius* | *collurio* | 3 | 4 | 0 | 0 | 7 | yes | |
| Pa | *Paridae* | *Parus* | *cristatus* | 2 | 5 | 0 | 0 | 7 | yes | |
| Pi | *Picidae* | *Dendrocopos* | *major* | 1 | 2 | 3 | 1 | 7 | yes | feed on insects[2)] |
| Pa | *Fringillidae* | *Carduelis* | *spinus* | 3 | 0 | 2 | 1 | 6 | yes | feed insects to the young |
| Pa | *Motacillidae* | *Anthus* | *pratensis* | 3 | 0 | 0 | 3 | 6 | yes | |
| Pa | *Fringillidae* | *Serinus* | *pusillus* | 0 | 0 | 6 | 0 | 6 | yes | feed insects to the young? |
| Pa | *Muscicapidae* | *Saxicola* | *rubicola* | 0 | 0 | 6 | 0 | 6 | yes | |
| Pa | *Corvidae* | *Corvus* | *corone* | 0 | 0 | 0 | 6 | 6 | yes | |
| Pa | *Muscicapidae* | *Ficedula* | *hypoleuca* | 3 | 2 | 0 | 0 | 5 | yes | |
| Pa | *Aegithalidae* | *Aegithalos* | *caudatus* | 0 | 5 | 0 | 0 | 5 | yes | |
| Pa | *Hirundinidae* | *Riparia* | *riparia* | 0 | 0 | 5 | 0 | 5 | no | hunt in the air |
| Pa | *Muscicapidae* | *Ficedula* | *parva* | 4 | 0 | 0 | 0 | 4 | yes | |
| Pa | *Muscicapidae* | *Saxicola* | *rubetra* | 2 | 0 | 1 | 1 | 4 | yes | |
| Pa | *Corvidae* | *Nucifraga* | *caryocatactes* | 0 | 4 | 0 | 0 | 4 | yes | |
| Pa | *Hirundinidae* | *Ptyonoprogne* | *rupestris* | 0 | 0 | 4 | 0 | 4 | no | hunt in the air |
| Pa | *Fringillidae* | *Carduelis* | *chloris* | 0 | 0 | 1 | 3 | 4 | yes | feed insects to the young |
| Pa | *Sittidae* | *Sitta* | *krueperi* | 0 | 0 | 3 | 0 | 3 | no | missing data |
| Pa | *Paridae* | *Parus* | *palustris* | 2 | 0 | 0 | 0 | 2 | yes | |
| Pa | *Muscicapidae* | *Phoenicurus* | *phoenicurus* | 2 | 0 | 0 | 0 | 2 | yes | |
| Pa | *Sylvidae* | *Acrocephalus* | *schoenobaenus* | 1 | 0 | 0 | 1 | 2 | yes | |
| Pa | *Corvidae* | *Pica* | *pica* | 0 | 1 | 0 | 1 | 2 | yes | |
| Pa | *Sylvidae* | *Phylloscopus* | *sindiatus l.* | 0 | 0 | 2 | 0 | 2 | yes | |
| Pa | *Hirundinidae* | *Hirundo* | *rustica* | 0 | 0 | 1 | 1 | 2 | no | hunt in the air |
| Pa | *Alaudidae* | *Alauda* | *arvensis* | 0 | 0 | 0 | 2 | 2 | no | observed in one transect only |
| Pa | *Fringillidae* | *Coccothraustes* | *coccothraustes* | 1 | 0 | 0 | 0 | 1 | yes | feed insects to the young |
| Pa | *Corvidae* | *Corvus* | *corax* | 1 | 0 | 0 | 0 | 1 | yes | |
| Pa | *Sylvidae* | *Locustella* | *fluviatilis* | 1 | 0 | 0 | 0 | 1 | yes | |
| Pa | *Turdidae* | *Turdus* | *iliacus* | 0 | 1 | 0 | 0 | 1 | yes | |
| Ch | *Scolopacidae* | *Tringa* | *ochropus* | 0 | 1 | 0 | 0 | 1 | no | |
| Pa | *Muscicapidae* | *Monticola* | *saxatilis* | 0 | 0 | 1 | 0 | 1 | yes | |
| Pa | *Muscicapidae* | *Phoenicurus* | *ochruros* | 0 | 0 | 1 | 0 | 1 | yes | |
| Pi | *Picidae* | *Picus* | *viridis* | 0 | 0 | 1 | 0 | 1 | yes | feed on insects |
| Pa | *Emberizidae* | *Emberiza* | *schoeniclus* | 0 | 0 | 0 | 1 | 1 | yes | feed insects to the young |

1) Milwright, R.D.P. (1998) Breeding biology of the golden oriole Oriolus oriolus in the fenland basin of eastern Britain. Bird Study, 45(3): 320-330
2) Leikola, Anto; Lokki, Juhani; Stjernberg, Torsten: Von Wright -veljesten linnut, s. 191. Otava, 2006. ISBN 951-1-18037-1

**Table S4 (related to Figure 3 in the main text).** Family level component loadings of the second and third principal components describing 33.7% and 8.5% of the total variation of bird communities across countries, respectively.

| Bird family | PC2 loading | PC3 loading |
|---|---|---|
| Motacillidae | 0.096621248 | 0.067106914 |
| Troglotydidae | 0.014708180 | 0.210943319 |
| Sylvidae | 0.009891431 | -0.430736201 |
| Oriolidae | 0.005022655 | -0.054848221 |
| Prunellidae | 0.001358338 | -0.042870544 |
| Corvidae | 0.001155818 | 0.011587582 |
| Picidae | -0.013150415 | -0.024347383 |
| Certhiidae | -0.014865169 | 0.011404222 |
| Turdidae | -0.019693893 | -0.007807002 |
| Laniidae | -0.022166815 | -0.032307565 |
| Muscicapidae | -0.032369494 | 0.532307129 |
| Fringillidae | -0.062877940 | 0.687239259 |
| Paridae | -0.991991538 | -0.054855828 |
